# *Brucella* targets host USP8 through the effector protein TcpB to facilitate infection of macrophages

**DOI:** 10.1101/2023.04.28.538663

**Authors:** Girish Radhakrishnan, Kiranmai Joshi, Varad Mujumdar

## Abstract

*Brucella* species are Gram-negative intracellular bacterial pathogens that cause the worldwide zoonotic disease brucellosis. *Brucella* can infect many mammals, including humans and domestic and wild animals. *Brucella* manipulates various host cellular processes to invade and multiply in professional and non-professional phagocytic cells. However, the host targets and their modulation by *Brucella* to facilitate the infection process remain obscure. Here, we report that the host Ubiquitin Specific Protease, USP8 negatively regulates the invasion of *Brucella* into macrophages through the plasma membrane receptor, CXCR4. *Brucella* suppressed the expression of USP8 at its early stage of infection in the infected macrophages. Subsequent studies revealed that the *Brucella* effector protein, TIR-domain containing protein from *Brucella*, TcpB plays a significant role in downregulating the expression of USP8 by targeting the CREB pathway. Treatment of mice with USP8 inhibitor resulted in enhanced survival of *B. melitensis,* whereas mice treated with CXCR4 or 14-3-3 antagonist showed a diminished bacterial load. Our experimental data demonstrate a novel role of USP8 in the host defence against microbial intrusion and microbial subversion of host defences.

## Introduction

Genus *Brucella* contains Gram-negative facultative intracellular bacterial pathogens, causing the worldwide zoonotic disease, brucellosis. *Brucella* can infect many mammals, including domestic and wild animals, marine mammals, and humans (Suárez-Esquivel et al., 2020). Based on the differences in pathogenicity and host preferences, twelve species of *Brucella* are currently recognized such as *B. melitensis* (goat and sheep), *B. abortus* (cattle), *B. suis* (swine), *B. neotomae* (desert rat), *B. ovis* (sheep), and *B. canis* (dogs), *B. pinnipedialis* (seals), *B. ceti* (cetaceans), *B. microti* (wood rat), *B. papionis* (baboons) and *B. inopinata* (humans) (Kurmanov et al., 2022). Brucellosis is highly contagious, and the primary cause of human infection is consuming contaminated dairy products, inhaling *Brucella*-containing aerosols, or contact with infected materials. Acute human infection manifests as generalized symptoms, including fever, chills, headache, and fatigue. The chronic infection may lead to endocarditis and meningitis, which can be fatal (Baldwin and Goenka, 2006). In animals, *Brucella* spp. cause abortions in the late trimester, stillbirths, retention of the placenta and decreased milk production in females and infertility in males, which is the major consequence leading to substantial economic loss in the livestock industry worldwide (Franc et al., 2018; Gheibi et al., 2018). Treatment of human *Brucellosis* involves prolonged oral regimens of doxycycline and streptomycin for 6 weeks, where 16% of individuals who undergo antibiotic treatment may show relapse (Skalsky et al., 2008). Frequent therapeutic failures also make antibiotic therapy for brucellosis ineffective. There is no human vaccine for brucellosis; the only option to control human infection is mass vaccination of susceptible animals. However, the available animal vaccines have significant drawbacks, including their infectivity to humans (Zai et al., 2021). Therefore, understanding the *Brucella*-host interaction is crucial to identify novel targets for developing improved vaccines and therapeutics for brucellosis.

*Brucella spp.* invade and multiply in macrophages, dendritic cells, trophoblasts, and epithelial cells (Salcedo et al., 2013). *Brucella* employs a secretion system to introduce bacterial effectors into the infected cells. These proteins interfere with host cellular pathways allowing the pathogen to resist intracellular killing and build an intracellular niche favourable for replication. *Brucella* harbours a Type IV Secretory System (T4SS) encoded by the VirB operon that is involved in the secretion of many effector proteins in the infected macrophages. These effector proteins interact with components of cellular pathways to generate replication-permissive, ER-derived compartments, leading to the chronic persistence of bacteria in the host (Celli et al., 2003). Since the interplay between bacterial effectors and the host cellular machinery plays a critical role in the invasion and persistence of *Brucella*, understanding these mechanisms is crucial for developing effective therapeutic and preventive measures for brucellosis. Studies have identified a few host proteins *Brucella* targets during their multistage intracellular cycle involving entry, trafficking, replication, and egress out of the host cells (Celli, 2019). Brucellae interact with GTPases belonging to the Rho subfamily, *viz*. Rho-Rac-CDC42 induces cytoskeleton remodeling during cell invasion (Guzmán-Verri et al., 2001). Many cell surface receptors such as CXCR4, CDC36, and PrpC are reported to be induced by *Brucella* infection, which promote entry into the macrophages (Nakato et al., 2012; Reyes et al., 2019). Small GTPases such as Sar1 and Rab2 were reported to play an essential role in the intracellular replication of *Brucella* (Celli et al., 2005; Smith et al., 2020). Brucellae are known to activate Unfolded Protein Response (UPR) by phosphorylating IRE1 through Yip1 to form a replicative niche in the endoplasmic reticulum (Li et al., 2022; Smith et al., 2013; Taguchi et al., 2015). They also target various components of host immune signalling pathways, including Toll-like receptors, to evade or suppress immune responses that contribute to their chronic persistence in the host (Alaidarous et al., 2014; Jakka et al., 2017; Radhakrishnan et al., 2009). However, evidence suggests that *Brucella* manipulates host cellular processes and signalling pathways for their survival, our understanding of the *Brucella*-host interaction is still rudimentary compared to other invasive bacterial pathogens.

By employing a siRNA-based screening, we identified that the host Ubiquitin Specific Protease-8 (USP8) plays a crucial role in *Brucella* infection of macrophages. USP8 is a promiscuous deubiquitinating enzyme that counterbalances the ubiquitination harnessed by numerous E3 ligases (De Ceuninck et al., 2013). It maintains the dynamic state of cellular ubiquitome by removing the conjugated ubiquitin moiety from the substrate proteins. USP8 selectively interacts with a palette of substrates through its substrate-binding motif and is involved in sorting out various membrane receptors for either recycling or degradation (Dufner and Knobeloch, 2019). USP8 is reported to regulate endosomal trafficking by stabilizing the endosomal sorting complex (ESCRT-0) through deubiquitinating the scaffold proteins Hrs and STAM1/2 (Berlin et al., 2010b). We observed that silencing of USP8 enhanced the uptake of *Brucella* into macrophages, whereas its overexpression suppressed the *Brucella* invasion. In addition, USP8 affected the interaction of *Brucella* with macrophages through the regulation of the availability of the plasma membrane receptor, CXCR4. Further, we found that *Brucella* suppressed the expression of USP8 at the initial stages of its infection through the effector protein, TcpB. The modulation of USP8 activity using the inhibitors or activators affected the splenic load of *B. melitensis* in the infected mice. Our research findings uncovered a novel role of the host protein, USP8, which plays a vital role in the defence against infectious disease. The study also signified the strategies employed by the infectious pathogens to counteract this host defence mechanism to establish the infection.

## Materials and methods

### Ethical statement

All the experiments involving *B. melitensis* were conducted at the Biosafety Level-3 /Animal -BSL3 facility of NIAB. The experimental protocols were approved by the Institutional Biosafety Committee (Approval number: IBSC/Jul2020/NIAB/GR01) and Institutional Animal Ethics Committee (Approval number: IAEC/NIAB/2022/10/GKR) and BSL-3 Research Review Committee (Approval number: BSL3-Jan2022/002). Six to eight-week-old female BALB/c mice were procured from the Small Animal Facility (SAF) of the National Institute of Animal Biotechnology (NIAB).

### Cell culture

Immortalized Bone Marrow-Derived Macrophages from Mice (iBMDMs; a gift from Petr Broz, University of Lausanne) and Human Embryonic Kidney (HEK) 293T cells (ATCC) were cultured in Dulbecco’s Modified Eagle’s Medium (DMEM; Sigma) supplemented with 10% fetal bovine serum (Sigma) and 1X penicillin-streptomycin solution (Gibco). RPMI 1640 medium (Sigma) supplemented with 10% fetal bovine serum and 1X penicillin-streptomycin solution was used to culture the murine macrophage cell line, J774 (ATCC). The cells were grown in a humidified atmosphere of 5% CO_2_ at 37° C. The iBMDMs were differentiated using m-CSF (Bio Legend; 20 ng/ml) or the culture supernatant of L929 cells (ATCC).

### Silencing and overexpression of USP8 in macrophages

To silence the USP8 gene in iBMDMs, a set of four siRNAs was procured from Dharmacon (ON-TARGETplus SMARTpool). The iBMDMs were seeded (5×10^4^ cells/well) into 24-well plates and transfected with 50 picomoles of siUSP8 or non-target (NT) siRNA in duplicates using Dharmafect 4.0 transfection reagent as per manufactures protocol. To examine the silencing of USP8 in iBMDMs, the cells were harvested 48 hours post-transfection, followed by total RNA isolation and cDNA synthesis. Subsequently, qRT-PCR analysis was performed using USP8-specific primers to quantify the mRNA expression levels of USP8. Data were normalized with the endogenous control, GAPDH. Further, the siRNA-transfected iBMDMs were subjected to immunoblotting using the anti-USP8 antibody to detect the endogenous levels of USP8. Actin was used as the loading control.

To overexpress USP8, iBMDMs were seeded (5×10^4^ cells/well) into 24-well plates and allowed to adhere overnight. Next, the cells were transfected with the eukaryotic expression plasmid harbouring USP8 (pCMV-HA-USP) or Empty vector (pCMV-HA) using X-fect (TAKARA Bio) transfection reagent according to the manufacturer’s instructions. The overexpression of USP8 in iBMDMs was confirmed by qRT-PCR and immunoblotting, as described before.

### Infection of macrophages with *B. neotomae* or *B. melitensis*

To perform infection studies using USP8-silenced iBMDMs, the cells were first transfected with siRNA. Forty-eight hours post-transfection, the cells were infected with *B. neotomae* (ATCC 23459-5K33) at an MOI of 1000:1 or *B. melitensis* 16M (obtained from Indian Veterinary Research Institute) at an MOI of 200:1. After the addition of *Brucella*, the plates were centrifuged at 280 X g for 3 min to pellet down the bacteria onto the cells.

Subsequently, the plates were incubated for 90 min at 37° C with 5% CO2. Next, the cells were washed three times with phosphate-buffered saline (PBS) and treated with gentamicin (30 µg/mL) for 30 min to kill extracellular *Brucella*. The infected cells were maintained in the cell culture media containing 3 µg/mL of gentamycin. Next, the cells were lysed with 0.1% Triton X-100 in PBS at various hours post-infection, followed by serial dilution of lysates and plating on brucella agar (BD). The intracellular load of *Brucella* was quantified by enumerating the CFU, and the data were represented as CFU/mL. To perform infection studies using USP8 overexpressing iBMDMs, the cells were infected with *B. neotomae* or *B. melitensis* 16M 24 hours post-transfection as described before.

For the invasion assay, *B. neotomae* or *B. melitensis* was added to the multi-well plates harbouring the macrophages, followed by centrifuging the bacteria onto the cells. Subsequently, the plates were incubated for 30 min at 37° C with 5% CO2. Next, the cells were washed thrice with PBS and treated with 30 µg/mL gentamicin for 30 min to kill extracellular *Brucella*. Subsequently, the infected cells were lysed with 0.1% Triton X-100 in PBS, then serial dilution of lysates and plating on brucella agar. Finally, the intracellular *Brucella* was quantified by enumerating the CFU. To perform invasion assay using USP8 overexpressing iBMDMs, the cells were transfected with the eukaryotic expression plasmid harbouring USP8 (pCMV-HA-USP) or Empty vector (pCMV-HA) as described previously. Twenty-four hours post transfection, the cells were infected with *B. neotomae* for invasion assay as described above.

To perform macrophage invasion assay with *B. neotomae* or *B. melitensis* in the presence of inhibitors of USP8 (DUB-IN-2; MedChem; 10 µM) or CXCR4 (AMD3100; Sigma 25 µg/ml) or CREB (CREB Inhibitor; Sigma, 5 µM), iBMDMs were treated with CREB inhibitor for 3 hours or USP8 and CXCR4 inhibitor for 24 hours, followed by infection with *B. neotomae* or *B. melitensis* for 30 minutes and CFU analysis. For analyzing the *Brucella* invasion in the presence of the USP8 activator (BV02, Sigma, 20 µM), iBMDMs were treated with BV02 for 24 hours, followed by *Brucella* invasion assay as described before. The vehicles, such as DMSO (for USP8, CREB inhibitor, and USP8 activator) and PBS (for CXCR4), were used as the controls.

To evaluate the endogenous protein levels of USP8 or CREB/p-CREB, iBMDMs were infected with either *B. neotomae* or *B. melitensis.* The cells were harvested at various times post-infection, followed by cell lysis in RIPA buffer. The lysates were then clarified, and the amount of total protein in the samples was quantified using the Bradford assay (Sigma). Subsequently, the samples were subjected to immunoblotting, and the levels of USP8 or CREB/p-CREB were detected using the respective primary antibodies and HRP-conjugated secondary antibodies (Table. 1).

To examine the expression of USP8 in the macrophages infected with heat-killed *Brucella*, *B. neotomae* was heat-inactivated by incubating the culture at 60° C for 1 hour in a water bath. Subsequently, the inactivation of *B. neotomae* was confirmed by streaking the culture on brucella agar plates. Next, iBMDMs were infected with heat-killed *B. neotomae*, and then the target gene expression was analyzed by qRT-PCR and immunoblotting.

### Immunoblotting

The harvested cells were lysed in RIPA buffer with a protease inhibitor cocktail (Pierce). The amount of total protein in the samples was quantified using the Bradford assay (Sigma). An equal concentration of protein samples was mixed with 2X Laemmli buffer (BioRad), and the samples were boiled for 10 min at 100° C. Next, the protein samples were resolved on SDS-PAGE gel, followed by the transfer of protein onto the PVDF membrane (Merck Millipore) using a wet-tank blotting system (Bio-Rad). The membrane was blocked with 5% skimmed milk in Tris-Buffered Saline with Tween 20 (TBST; Cell Signaling Technology) for 1 h, followed by incubation with the respective primary antibody overnight at 4° C. Next, the membrane was washed three times with TBST and incubated with horseradish peroxidase (HRP)-conjugated secondary antibody. Next, the primary or secondary antibody was diluted in 5% skimmed milk in TBST. Finally, the membrane was washed three times with TBST and incubated with Super Signal West Pico or Femto chemiluminescent substrate (Pierce). The signals were captured using a chemi-documentation system (Syngene). The antibody source and dilutions used are listed in Table.1.

**Table.1.** Details of antibodies used for the study

| S.No | Antibody | Manufacturer | Dilution |
| --- | --- | --- | --- |
| 1. | Anti-FLAG HRP-conjugated | Sigma | 1:5000 |
| 2. | Anti-HA HRP-conjugated | Sigma | 1:5000 |
| 3. | Anti-actin HRP-conjugated | Sigma | 1:25000 |
| 4. | Anti-USP8 | Thermo Fisher Scientific | 1: 1000 |
| 5. | Anti-CXCR4 | Invitrogen | 1: 1000 (WB),<br>1:500 (IF) |
| 6. | Anti-phospho-CREB | Cell Signalling Technology | 1: 1000 |
| 7. | Anti-CREB | Cell Signalling Technology | 1: 1000 |
| 8. | Anti-rabbit HRP-conjugated | Cell Signalling Technology | 1:5000 |
| 9. | Anti-mouse HRP-conjugated | Cell Signalling Technology | 1:5000 |
| 10. | Anti-calreticulin | Invitrogen | 1:200 |
| 11. | Anti-rabbit IgG Alexa Fluor<br>647 | Cell Signalling Technology | 1:500 |
| 12. | Anti-HA-FITC antibody | Sigma | 1:500 |
| 13. | Anti-TNFR1 | R&D | 1:500 |

### Gene expression analysis using quantitative RT-PCR

To analyse the expression of USP8 in the *Brucella*-infected cells by qRT-PCR, iBMDMs were seeded into 24-well plates (5×10^4^ cells/well) in DMEM supplemented with 10% FBS without antibiotics and incubated overnight at 37° C with 5% CO2. The cells were infected with *B. neotomae* (live or heat-killed) at MOI of 1000:1 or *B. melitensis* at MOI of 200:1 as described before, followed by harvesting the cells at various time points. The total RNA was isolated from the infected macrophages using RNAiso PLUS (Takara Bio), followed by the preparation of cDNA using Prime Script™ RT Reagent kit (Takara Bio) as per the manufacturer’s protocol. The qRT-PCR was performed using gene-specific primers with the SYBR Green method using the real-time PCR machine (BioRad). The relative gene expression was analysed by the comparative 2^-ΔΔCt^ method using CFX 96 software (BioRad). Data were normalized with the endogenous control, GAPDH. To examine the downregulation of target gene expression by siRNA, the cells were harvested 48 hours post-transfection, followed by RNA isolation, cDNA preparation, and qRT-PCR analysis. The overexpression of USP8 in macrophages was also confirmed by qRT-PCR analysis, as described before. All the primers used for analysing the gene expression by qRT-PCR are listed in Table 2. To assess the levels of USP8 in the macrophages overexpressing TcpB, iBMDMs were seeded into 24-well plates (5×10^4^ cells/well) and transfected with 600 ng and 1200 ng of the eukaryotic expression vector (pCMV-HA; Clontech) harbouring TcpB for 24-hours. Subsequently, the cells were collected and processed for qRT-PCR and immunoblotting as described before.

**Table.2:**
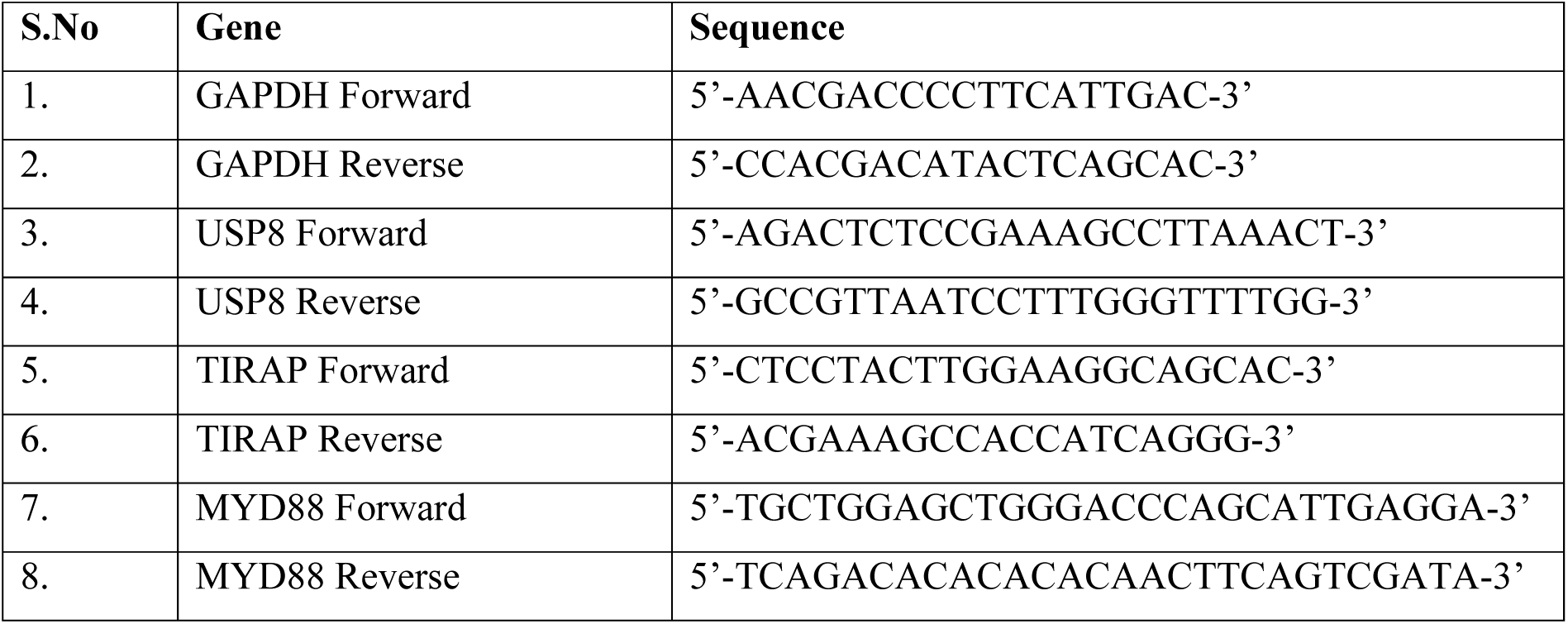
Primer sequences of genes used for qRT-PCR analysis.

### Preparation of membrane fraction

The total membrane fraction containing the CXCR4 protein was isolated from iBMDMs, as mentioned elsewhere (Yaseen et al., 2018). Briefly, iBMDMs were sonicated in 100 μl buffer containing 100 mM Tris–HCl pH 10.7, 5 mM EDTA, and 2 mM DTT. After sonication, the sample was diluted to 500 μl with buffer containing 100 mM Tris–HCl pH 8, 0.33 M sucrose, 5 mM EDTA, and 2 mM DTT. Next, the sample was centrifuged for 3 min at 1,000 *g*. Then, the supernatant was collected and centrifuged again for 5 min at 3,000 *g*. Finally, the resulting supernatant was centrifuged at 19,000 *g* for 45 min, followed by the collection of pellets corresponding to the total membrane fraction. The pellet was then dissolved in 50 μl buffer containing 10 mM Tris–HCl pH 7.5, 0.1 mM DTT, and 20% glycerol for further experiments.

### Immunofluorescence microscopy

To generate *B. neotomae* expressing GFP, the *Brucella* expression plasmid harboring GFP (pNSTrcd-GFP) was electroporated into *B. neotomae* using a MicroPulser (Bio-Rad). The transformants were selected on brucella agar containing chloramphenicol (40 µg/mL). To perform confocal microscopy analysis, macrophages infected with *B. neotomae*-GFP were washed with PBS and fixed with 4% paraformaldehyde for 20 min. The cells were then permeabilized with 0.1% Triton X-100 in PBS for 10 min. Subsequently, the cells were blocked with 1% BSA in PBS containing 50 mM NH_4_Cl for 30 min. The cells were then incubated with the anti-calreticulin antibody for 1 h, followed by Alexa Fluor 647-conjugated secondary antibody to stain the endoplasmic reticulum. Finally, the cells were mounted in Prolong Gold antifading agent with DAPI (Thermo Fisher Scientific). The cells were analyzed using a laser confocal microscope at 40X magnification (Leica). To perform fluorescence microscopy, the *B. neotomae*-GFP infected cells were fixed with 4% paraformaldehyde in PBS, followed by staining of the nucleus with Hoechst stain (Thermo Fisher Scientific). The images were captured using a fluorescent microscope (Carl Zeiss) at 20X magnification. Fifteen fields were analyzed from the samples transfected with NT and siUSP8 and quantified using Image J software.

To perform immunofluorescence microscopy for analysing membrane localization of CXCR4 in USP8 overexpressing cells, HEK 293T cells were seeded onto glass bottom dishes (0.5 x 10^6^ /Dish) in DMEM supplemented with 10% FBS and allowed to adhere overnight at 37° C with 5% CO2. Cells were then transfected with eukaryotic expression plasmid harbouring USP8 (pCMV-HA-USP8) or Empty vector using the X-fect transfection reagent. Twenty-four hours post-transfection, cells were washed with PBS and fixed with 4% paraformaldehyde for 20 minutes, followed by permeabilization with 0.1% Triton X 100 in PBS. Cells were then blocked with 1% BSA in PBS containing 50 mM NH_4_Cl. Next, the cells were stained with the anti-CXCR4 antibody, followed by AlexaFluor-647 secondary antibody, and HA-USP8 was stained with mouse anti-HA-FITC antibody (Table. 1). Finally, the cells were mounted in Prolong Gold anti-fading agent with DAPI. The cells were analysed using a fluorescence microscope (Carl Zeiss) at 20X magnification. To analyse the membrane localization of CXCR4 with USP8 inhibitor, iBMDMs were treated with DMSO or DUB-IN-2 (10 μM) for 24 hours. Subsequently, the cells were stained for CXCR4 as described above and analyzed using a laser confocal microscope (Leica) at 63X magnification. To analyse the membrane localization of CXCR4 with USP8 inhibitor, iBMDMs were treated with DMSO or DUB-IN-2 (10 μM) for 24 hours. Subsequently, the cells were processed for confocal microscopy as described above. Twelve different fields were analyzed from DUB-IN-2 or DMSO-treated samples.

To examine the invasion of *Brucella* in USP8 overexpressing cells, iBMDMs were transfected with pCMV-HA-USP8 or Empty vector using the X-fect transfection reagent. Twenty-four-hour post-transfections, the cells were infected with *B. neotomae-*GFP, followed by processing the cells for fluorescent microscopy as described before. The cells were analysed using a fluorescence microscope (Carl Zeiss) at 20X magnification, and *B. neotomae-*GFP per well was quantified using ImageJ software. Twelve different fields were analyzed for each sample.

### Cytotoxicity assay

To examine the cytotoxicity of DUB_IN-2 (USP8 inhibitor) and BV02 (14-3-3 inhibitor), iBMDMs were seeded into a 48-well plate and allowed to adhere overnight. The cells were treated with 5, 10, 15, and 20 µM of either DUB_IN-2/BV02 or DMSO for 24 hours. The supernatants were collected, then the released lactate dehydrogenase (LDH) level was quantified using CytoTox 96® (Promega) as per the manufacturer’s protocol. The untreated cells lysed with 0.1% Triton X100 served as the high control for the LDH assay.

### Co-transfection and protein degradation experiments

To examine the role of *Brucella* effector protein, TcpB, on negatively regulating the expression of USP8, HEK293T cells were transfected with eukaryotic expression plasmid harbouring TcpB. HEK293T cells were seeded into 12-well plates (1×10^5^ cells/well) and transfected with increasing concentrations (0, 600, 1200 ng) of TcpB expression plasmid. Twenty-four hours post-transfection, the cells were harvested and processed for immunoblotting as described previously. The blots were probed with HRP-conjugated anti-HA to detect TcpB, and the endogenous level of USP8 was detected using the anti-USP8 antibody, followed by HRP-conjugated anti-mouse IgG. To examine the mRNA expression of USP8, HEK293T cells were transfected with the expression plasmid harbouring TcpB. Twenty-four hours post-transfection, the cells were harvested and processed for RNA isolation and cDNA synthesis. Subsequently, qRT-PCR was performed to quantify the endogenous levels of USP8, TIRAP, and MYD88.

To examine the endogenous levels of TIRAP, p-CREB, and USP8 in the presence of TcpB, HEK 293T cells were seeded in 12-well (1×10^5^ cells/well) plates and transfected with various concentrations (0, 600, and 1200 ng) of pCMV-HA-TcpB. Twenty-four hours post-transfection, cells were harvested and processed for immunoblotting to examine the protein levels as described earlier. To determine whether the purified MBP-TcpB protein affects USP8 levels, iBMDMs were treated with purified MBP-TcpB protein or MBP alone (5 µg/ml) for 5 hours, followed by harvesting the cells and immunoblotting to detect the endogenous levels of USP8 protein. To analyse the degradation of TIRAP by TcpB, HEK293T cells were co-transfected with FLAG-TIRAP (300 ng) with increasing concentrations of HA-TcpB (0, 600, and 1200 ng). Twenty-four hours post-transfection, cells were harvested and subjected to immunoblotting to detect the expression levels of FLAG-TIRAP and HA-TcpB.

To examine whether overexpression of TIRAP leads to stabilization of USP8, HEK293T cells were transfected with pCMV-FLAG-TIRAP (1.2 µg). Twenty-four hours post-transfection, cells were harvested and processed for immunoblotting to assess the levels of phospho-CREB and USP8. For titrating the effect of TcpB on TIRAP-mediated expression of USP8, HEK293T cells were transfected with pCMV-HA-TcpB (600 ng), and two concentrations of pCMV-FLAG-TIRAP (600 and 1200 ng) for 24 hours. Subsequently, the cells were harvested and subjected to immunoblotting to detect the endogenous levels of USP8. Primary antibodies used for detecting TIRAP and p-CREB are listed in Table 1. The HRP-conjugated anti-rabbit and anti-mouse secondary antibodies were used to detect TIRAP, p-CREB, and USP8, respectively.

### Construction of TcpB knock-out *B. neotomae*

A homologous recombination-based gene replacement technique was used for deleting the TcpB gene from *B. neotomae*. A 3-way ligation was performed to generate the TcpB KO plasmid. The 1 kb fragments, upstream and downstream of the TcpB gene, were amplified from the chromosomal DNA of *B. neotomae*. The forward and reverse primers for amplifying the upstream fragment harboured *Kpn*I and *Bam*HI restriction enzymes, respectively. The *Bam*HI and *Eco*RI sites were added in the forward and reverse primers, respectively, to amplify the downstream fragment. The Kanamycin expression cassette from the pUC-kan-loxp plasmid was released with *Bam*HI digestion. The pZErO-1.1 plasmid (Invitrogen) was digested with *Kpn*I and *Xho1*, followed by ligating the upstream fragment with *Kpn*I & *Bam*HI, Kanamycin cassette with *Bam*HI and the downstream fragment with *Bam*HI & *Eco*RI restriction sites. The plasmid was transformed into DH5α, and the positive clones were confirmed by selecting on Zeocin and Kanamycin plates. Subsequently, the TcpB KO plasmid was introduced into *B. neotomae* by electroporation. The upstream and downstream fragments in the KO plasmid recombine with the respective sequences on the chromosomal DNA of *B. neotomae,* replacing the TcpB gene with the Kanamycin expression cassette. The transformed *B. neotomae* colonies growing on Kanamycin and Zeocin, which indicates a single recombination event and the resulting insertion of the KO plasmid into the chromosomal DNA, were discarded. The Zeocin sensitive and Kanamycin resistant TcpB KO *B. neotomae* colonies were selected and confirmed further by PCR.

To express TcpB in the *TcpB* KO *B. neotomae* for complementation experiments, the TcpB gene was cloned into the *Brucella* expression plasmid, pNSTrcD, at the *Sal*I and *Xho*I sites. *B. neotomae* harbouring pNSTrcD-TcpB was selected on brucella agar plates with chloramphenicol (40 µg/ml).

### *In-vivo* studies on mice using *B. melitensis*

To study the *in-vivo* effect of chemical inhibition or activation of USP8 or inhibition of CXCR4 in the mice model of brucellosis, 8-week-old female BALB/c mice (4 mice per group) were infected intraperitoneally with *B. melitensis* (2.5×10^6^ CFU per mouse) in 100 µL of 1X PBS. Ten days post-infection, mice were treated intraperitoneally with DUB_IN-2 or BV02, AMD3100, or vehicle control for 3 days. Mice were euthanized by CO2 asphyxiation on day 14, followed by the collection of spleens and CFU analysis to quantify the load of *B. melitensis* in the spleen.

### Statistical analysis

GraphPad Prism 6.0 software was used for the statistical analysis of experimental data. Data are shown as the mean standard ± error of the mean (SEM). Statistical significance was determined by t-tests (two-tailed) for pairwise comparison. A one-way analysis of variance (ANOVA) test was used to analyze the data involving more than two samples.

## Results

### USP8 plays an essential role in the *Brucella*-macrophage interaction

We identified the role of USP8 in the *Brucella-*macrophage interaction while performing a siRNA screening using *B. neotomae*, which is reported to be the model pathogen to study brucellosis (Kang et al., 2019) (Kang et al., 2019). The expression of USP8 in iBMDMs was downregulated by treating the cells with USP8-specific siRNA. The iBMDMs treated with siUSP8 exhibited a diminished expression of USP8, confirmed by qRT-PCR and immunoblotting (Figures. 1A & B). We observed an enhanced intracellular load of *B. neotomae* in iBMDMs treated with siUSP8 compared to the cells transfected with non-targeting (NT) control siRNA (Figure. 1C). Therefore, we wished to perform detailed studies to understand the role of USP8 in the *Brucella*-macrophage interaction. To study the effect of USP8 on the intracellular survival of *Brucella*, iBMDMs were treated with siUSP8 or NT, followed by infection with *B. neotomae* and analysis of CFU at various times post-infection. The CFU analysis indicated that the intracellular load of *B. neotomae* was significantly enhanced in the USP8-silenced macrophages at various times post-infection (Figure. 1D).

**Figure 1.**
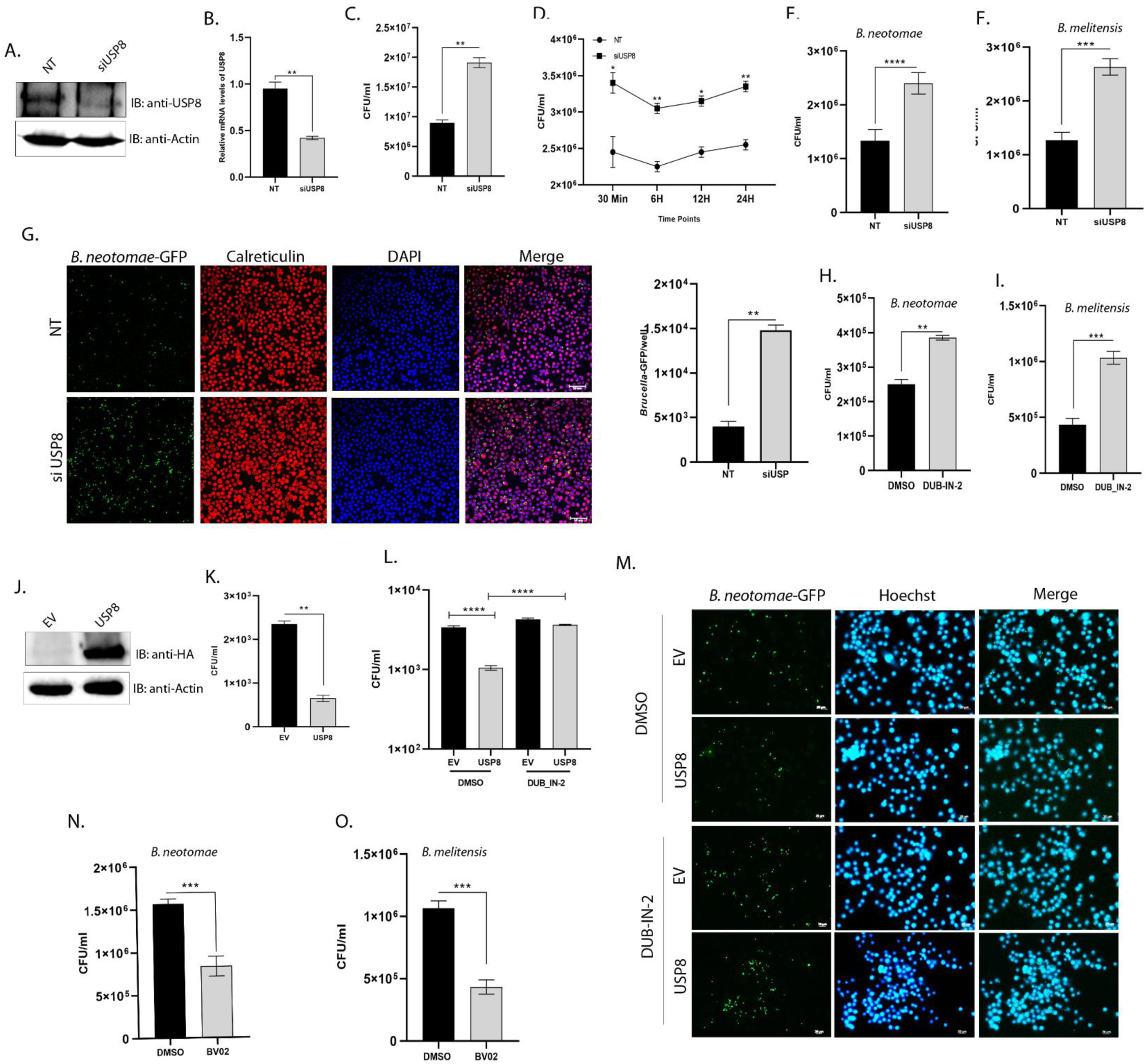
USP8 plays an essential role in *Brucella*-macrophage interaction. (A & B) Levels of endogenous USP8 treated with siUSP8 or non-targeting control (NT). The cells were treated with siRNAs for 48 hours, followed by analyzing the levels of USP8 by immunoblotting (A) and q-RT-PCR (B). (C & D) CFU analysis of *B. neotomae* isolated from USP8-silenced or control iBMDMs. The cells were infected with *B. neotomae*, followed by isolation of bacteria at 24 hours post-infection (C) or at indicated times (D) post-infection. (E & F) *Brucella* invasion assay using iBMDMs downregulating USP8 expression. iBMDMs were treated with siUSP8 or NT, followed by infection with *B. neotomae* or *B. melitensis* for 30 minutes. The cells were then treated with gentamicin for 30 minutes to kill extracellular bacteria, followed by quantification of invaded *B. neotomae* (E) or *B. melitensis* (F) by CFU enumeration. (G) *Brucella* invasion assay, followed by analyzing the invaded *B. neotomae*-GFP by confocal microscopy. The cells were stained with anti-calreticulin and Alexa Fluor 647-conjugated secondary antibody to visualize the endoplasmic reticulum (red). The nuclei were stained with DAPI (blue), which was present in the mounting reagent. Scale bar, 20 µm. The right panel indicates the quantification of intracellular *B. neotomae*-GFP using Harmony high-content analysis software. (H & I) *Brucella* invasion assay in the presence of USP8 inhibitor. iBMDMs were treated with DUB_IN-2 or DMSO (vehicle) for 24 hours, followed by infection with *B. neotomae* (H) or *B. melitensis* (I) and CFU analysis. (J) Immunoblot showing the overexpression of USP8 in iBMDMs. Cells were transfected with the eukaryotic expression plasmid harboring HA-tagged USP8 or empty vector. Twenty-four hours post-transfection, the cells were lysed, and the lysates were subjected to immunoblotting. The membrane was probed with HRP-conjugated anti-HA antibody to detect the overexpressed HA-USP8. (K) *Brucella* invasion assay using iBMDMs overexpressing HA-USP8. iBMDMs were transfected with HA-USP8 expression plasmid, followed by infection with *B. neotomae* and quantification of invaded bacteria by CFU enumeration. (L & M) *Brucella* invasion assay using iBMDMs overexpressing HA-USP8 in the presence or absence of USP8 inhibitor. iBMDMs overexpressing USP8 were treated with DUB_IN-2 or DMSO, followed by infection with *B. neotomae* or *B. neotomae*-GFP. The invasion of *B. neotomae* and *B. neotomae*-GFP were examined by CFU enumeration (L) and fluorescence microscopy (M), respectively. (N & O) *Brucella* invasion assay using iBMDMs treated with 14-3-3 inhibitor, BV02. The cells were treated with BV-02 or DMSO for 24 hours, followed by infection with *B. neotomae* (N) or *B. melitensis* (O) and CFU analysis. The data shown in A, B, C, D, J, K, L & M panels are representations of two independent experiments. The data shown in E, F, G, H, I, N & O panels are representations of three independent experiments. The data are presented as the mean ± SEM (P < 0.05*; P < 0.01**, P <0.001; ***).

Since we observed a high bacterial load even at the early stages of infection, we performed an invasion assay to examine whether USP8 plays any role in the uptake of *Brucella* by macrophages. iBMDMs were treated with siUSP8 or NT, followed by infection with *B. neotomae* or *B. neotomae*-GFP for 30 minutes and treatment with gentamycin to kill the extracellular *Brucella.* The CFU analysis showed a remarkable enhancement of *B. neotomae* invasion in the USP8-silenced cells compared to the control (Figure. 1E). Next, we infected USP8-silenced macrophages with the zoonotic species of *Brucella*, *B. melitensis,* which also showed an enhanced invasion in the USP8-downregulated iBMDMs (Figure. 1F). Similarly, the fluorescent microscopy analysis showed an enhanced number of *B. neotomae*-GFP in the siUSP8-treated iBMDMs compared to the control (Figure. 1G). Further, to validate the role of USP8 on the invasion of *Brucella*, we inhibited USP8 using the compound, DUB-IN-2, followed by infection studies using *B. neotomae* or *B. melitensis*. DUB-IN-2 was reported to inhibit the USP8, and treatment of iBMDMs with DUB-IN-2 for 24 hours neither induced any cytotoxicity nor modulated the expression of USP8 (Kageyama et al., 2020) (Figures. S1A & B). Next, the inhibitor or the vehicle (DMSO) treated iBMDMs were infected with *B. neotomae* or *B. melitensis*, followed by the CFU analysis. In agreement with the silencing data, chemical inhibition of USP8 also showed an enhanced invasion of both *B. neotomae* and *B. melitensis* into macrophages (Figures. 1 H & I).

To further confirm the experimental data, we overexpressed USP8 in the murine macrophages, then analysed the invasion of *B. neotomae*. To overexpress USP8, the iBMDMs were transfected with the eukaryotic expression plasmid harbouring HA-tagged USP8 or empty vector for 24 hours. The overexpression of USP8 in the transfected iBMDMs was confirmed by q-RT-PCR analysis and immunoblotting (Figure. S1C, Figure. 1J). Next, the iBMDMs overexpressing USP8 were infected with *B. neotomae* or *B. neotomae*-GFP for the invasion assay. The CFU analysis showed a significant reduction in the invasion of *B. neotomae* in USP8 overexpressing macrophages compared to the cells transfected with the empty vector (Figure. 1K). Next, we wished to examine whether the effect of USP8 overexpression is negated by inhibiting its function using the antagonist. The iBMDMs overexpressing USP8 were treated with DUB-IN-2 for 24 hours, followed by infection with *B. neotomae*/*B. neotomae*-GFP. We observed an enhanced intracellular load of *B. neotomae*/*B. neotomae*-GFP in USP8-overexpressing cells that are treated with DUB-IN-2 as compared to cells treated with DMSO (Figures. 1L & M). The protein 14-3-3 is reported to bind to USP8, which negatively regulates the activity of USP8 (Centorrino et al., 2018; Mizuno et al., 2007). Therefore, inhibition of USP8 and 14-3-3 interaction can lead to the constitute activation of USP8. We used the commercially available peptide, BV2, that interferes with the interaction of USP8 with 14-3-3, followed by *Brucella* infection studies. The iBMDMs were treated with BV2 or DMSO for 24 hrs and infected with *B. neotomae* or *B. melitensis*. We confirmed that BV2 did not induce any cytotoxicity of iBMDMs upon treatment for 24 hours (Figure. S1D). In agreement with the overexpression studies, we observed a diminished invasion of *B. neotomae* or *B. melitensis* into iBMDMs treated with BV02 (Figures. 1N & O). Collectively our experimental data show that USP8 negatively regulates the invasion of *Brucella* into macrophages.

### USP8 affects the invasion of *Brucella* into macrophages through the CXCR4 receptor

Since USP8 affected the invasion of *Brucella* into macrophages, we wished to examine whether USP8 regulates the membrane receptors required for *Brucella* entry. USP8 is known to negatively regulate the membrane localization and turnover of CXCR4, which is reported to be an essential receptor for the entry of *Brucella* into macrophages (Berlin et al., 2010a; Reyes et al., 2019). Since overexpression of USP8 inhibited the invasion of *Brucella* into macrophages, we hypothesized that USP8 might exert this effect by depleting the membrane-localized CXCR4 through negative regulation of its turnover inside the cells by promoting proteasomal degradation. To examine this, we analysed levels of plasma membrane-localized CXCR4 in iBMDMs transfected with USP8 expression plasmid or empty vector. Twenty-four hours post-transfection, the cells were fixed and stained for HA-USP8 and endogenous CXCR4, followed by fluorescence microscopy analysis. We observed a decreased membrane localized CXCR4 in the cells overexpressing USP8 compared to those transfected with the empty vector (Figure. 2A). Further, to confirm whether USP8 can regulate the turnover of CXCR4, we examined the total and membrane-localized CXCR4 in iBMDMs treated with USP8 inhibitor, DUB-IN-2 or USP8 activator, BV02. The immunoblotting showed an enhanced level of CXCR4 in total cell lysates and the membrane fractions upon treatment with the USP8 inhibitor. In contrast, the levels of CXCR4 decreased in the cells treated with the USP8 activator, indicating the role of USP8 in regulating the turnover of CXCR4 (Figures. 2B, C, D & E). The overexpression of USP8 in iBMDMs also resulted in a diminished level of CXCR4 as compared to the empty vector (Figure. S2A). Further, we examined the membrane localization of CXCR4 in the *Brucella*-infected macrophages. The iBMDMs were infected with *B. neotomae* for 4 hours, followed by isolation of plasma membrane fraction and immunoblotting. An enhanced membrane localized CXCR4 was detected in iBMDMs infected with *B. neotomae* compared to the uninfected cells (Figure. 2F). Our fluorescent microscopy studies also showed an enhanced level of CXCR4 on the plasma membrane in iBMDMs that are treated with USP8 inhibitor or infected with *B. neotomae* as compared to the controls (Figure. 2G & Figure. S2B)

**Figure 2.**
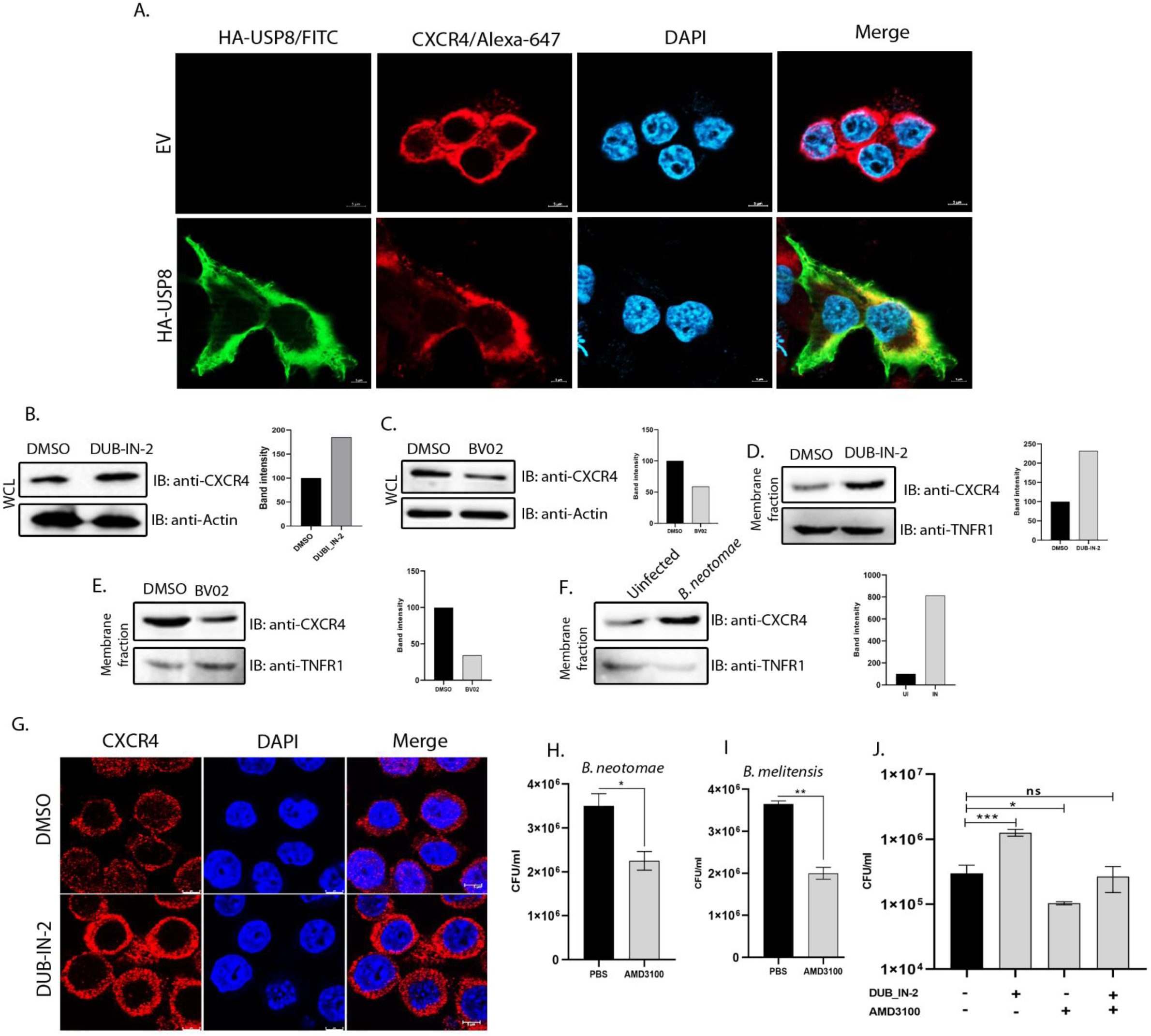
USP8 affects the invasion of *Brucella* into macrophages through CXCR4. (A) Levels of CXCR4 on the plasma membrane of HEK293T cells overexpressing HA-USP8. HEK293T cells were transfected with eukaryotic expression plasmid harboring HA-USP8 or empty vector. Twenty-four hours post-transfection, the cells were stained with the anti-CXCR4 antibody, followed by Alexa Fluor 647-conjugated secondary antibody (red). HA-USP8 was stained with FITC-conjugated anti-HA antibody (green), and nuclei were stained with DAPI (blue). The cells were imaged using a laser confocal microscope at 63X. Scale bar, 5μm. The image represents twelve different fields captured from cells transfected with either empty vector or HA-USP8. (B & C) Total levels of CXCR4 in iBMDMs treated with antagonists of USP8 or 14-3-3. iBMDMs treated with DUB_IN-2 (B) or BV02 (C), or DMSO (vehicle) for 24 hours, followed by harvesting the cells and immunoblotting. To detect CXCR4, the membranes were probed with the anti-CXCR4 antibody, followed by HRP-conjugated anti-rabbit IgG. Actin was used as the loading control. The right panels indicate the densitometry of the CXCR4 band with respect to the actin band. (D-F) Levels of plasma membrane-localized CXCR4 from iBMDMs treated with DUB-IN-2 (D) or BV02 (E) or infected with *B. neotomae* (F). The membrane fractions were isolated from the compound-treated or *Brucella*-infected cells, followed by immunoblotting to detect CXCR4. TNFR1 was used as the positive control, detected using the anti-TNFR1 antibody, followed by HRP-conjugated anti-mouse IgG. The right panel indicates the densitometry of the CXCR4 band with respect to the TNFR1 band. (G) iBMDMs showing membrane localized CXCR4 upon treatment with DUB_IN-2 or DMSO. iBMDMs were treated with DUB_IN-2 or DMSO for 24 hours, followed by staining membrane localized CXCR4 (red) as described earlier. Nuclei were stained with DAPI (blue). The cells were imaged using a laser confocal microscope at 63X magnification. Scale bar, 20 μm. The image represents twelve different fields captured from cells treated with either DMSO or DUB-IN-2. (H & I) *Brucella* invasion assay in the presence of the CXCR4 inhibitor. iBMDMs were treated with AMD3100 or PBS (vehicle) for 24 hours, followed by infection with *B. neotomae* or *B. melitensis* for 30 minutes. The cells were then treated with gentamicin to kill extracellular bacteria for 30 minutes, followed by quantification of invaded *B. neotomae* (H) or *B. melitensis* (I) by CFU enumeration. (J) *Brucella* invasion assay using iBMDMs treated with antagonists of USP8 and CXCR4, as indicated in the figure. iBMDMs were treated with DUB_IN-2 or AMD3100 or both DUB_IN-2 and AMD3100 for 24 hours, followed by infection with *B. neotomae* and enumeration of CFU. The data shown in A, G, H, I & J panels are representations of two independent experiments. The data shown in B, C, D, E & F panels are representations of three independent experiments. The data are presented as the mean ± SEM (P < 0.05*; P < 0.01**, P <0.001; ***). WCL: Whole cell lysate.

Subsequently, we analysed the role of CXCR4 on the invasion of *Brucella* into the macrophages. iBMDMs were treated with the antagonist of CXCR4, AMD3100, or vehicle control for 24 hours, followed by analysing the invasion of *B. neotomae* or *B. melitensis*. We observed a significant decline in the entry of *Brucella* into the cells that are treated with AMD3100 as compared to the vehicle-treated cells, suggesting that the CXCR4 receptor plays an essential role in the entry of *Brucella* into the macrophages (Figures. 2H & I). Further, to link the role of USP8 and CXCR4 in *Brucella* invasion, we examined the combined effect of USP8 and CXCR4 antagonists upon entry of *Brucella* into macrophages. The iBMDMs were treated with DUB-IN-2 or AMD3100 or both the antagonists, followed by invasion assay. As anticipated, the higher *Brucella* invasion observed in macrophages treated with USP8 inhibitor was negated by treatment with CXCR4 inhibitor (Figure. 2J). Our experimental data suggest that CXCR4 acts as an active receptor for *Brucella* invasion, and USP8 negatively regulates the plasma membrane localization of CXCR4, which interferes with the invasion of *Brucella* into macrophages.

### *Brucella* modulates the expression of USP8 in macrophages

Since USP8 negatively regulated the invasion of *Brucella* into macrophages, we speculated that *Brucella* might modulate the expression of USP8 to overcome its impact on infecting the macrophages. Therefore, we examined the endogenous levels of USP8 in the macrophages infected with *B. neotomae* or *B. melitensis* at various times post-infection. We observed a significant suppression of mRNA expression of USP8 at the early time points up to 4 hours and restoration of USP8 levels by 8 hours in the macrophages infected with *B. neotomae* or *B. melitensis* (Figures. 3A & B). To confirm the experimental data, we examined the endogenous level of USP8 by immunoblotting in the *B. neotomae*-infected macrophages. In agreement with the mRNA expression data, the protein level of USP8 was also decreased at the early time points, and the expression was restored by 8 hours post-infection (Figure. 3C). Next, we sought to examine whether *Brucella* actively modulates the expression of USP8 in the infected macrophages. To examine this, iBMDMs were infected with heat-killed *B. neotomae*, followed by analysing the USP8 expression level by q-RT-PCR and immunoblotting. Interestingly, heat-killed *B. neotomae* did not modulate the expression of USP8, indicating the requirement of live *Brucella* for suppressing the expression of USP8 (Figures. 3D & E).

**Figure 3.**
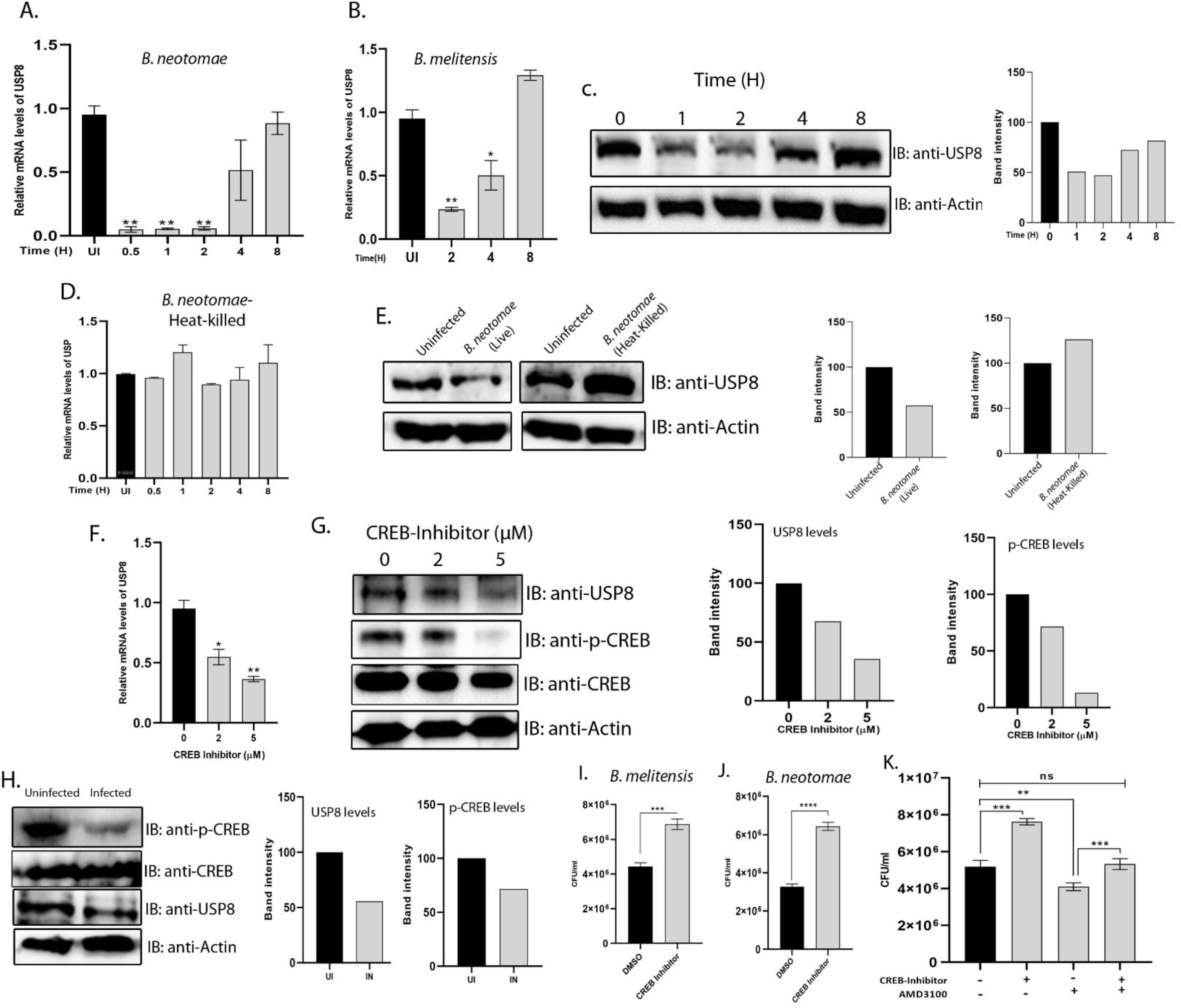
*Brucella* modulates the expression of USP8 in macrophages. (A & B) Endogenous levels of USP8 in iBMDMs infected with *Brucella*. iBMDMs were infected with *B. neotomae* (A) or *B. melitensis* (B), followed by harvesting the cells at indicated time points and quantifying USP8 mRNA expression by qRT-PCR. The expression of USP8 was normalized with the internal control, GAPDH, and the relative mRNA expression was determined compared to the uninfected cells. (C) Immunoblot showing USP8 levels in the *B. neotomae*-infected iBMDMs. The cells were infected with *B. neotomae* for the indicated time points, followed by cell lysis and immunoblotting. To detect endogenous USP8, the membranes were probed with the anti-USP8 antibody, followed by HRP-conjugated anti-mouse IgG. Actin was used as the loading control. The right panel indicates the densitometry analysis of USP8 bands normalized with the actin bands. (D) The endogenous level of USP8 in the iBMDMs infected with heat-killed *B. neotomae.* The cells were infected with heat-killed *B. neotomae* for the indicated time points, followed by quantification of USP8 level by qRT-PCR. (E) Immunoblot showing USP8 levels in iBMDMs infected with live or heat-killed *B. neotomae.* The cells were subjected to immunoblotting 4 hours post-infection, followed by detection of endogenous levels of USP8. The right panel indicates the densitometry analysis of USP8 bands normalized with the actin bands. (F & G) The endogenous levels of USP8 in iBMDMs treated with CREB inhibitor. The cells were treated with indicated concentrations of CREB inhibitor for 3 hours, followed by harvesting the cells. The cells were subjected to qRT-PCR analysis to determine the mRNA expression of USP8 (F) or immunoblotting to detect the protein levels of USP8 and phosphorylated-CREB (G). To detect the phosphorylated form of CREB, the anti-phospho-CREB antibody, followed by HRP-conjugated anti-rabbit IgG, was used. Actin was used as the loading control. The right panel indicates the densitometry analysis of USP8 and p-CREB bands normalized with actin bands. (H) Immunoblot showing USP8 and p-CREB in iBMDMs infected *B. neotomae.* The cells were infected for 4 hours, followed by cell lysis and immunoblotting to detect the endogenous levels of USP8 and phospho-CREB. The right panel indicates the densitometry analysis of USP8 and p-CREB bands normalized with actin bands. (I & J) *Brucella* invasion assay in the presence of CREB inhibitor. iBMDMs were treated with CREB Inhibitor or DMSO (vehicle) for 3 hours, followed by infection with *B. neotomae* or *B. melitensis* for 30 minutes. The cells were then treated with gentamicin to kill extracellular bacteria for 30 minutes, followed by quantification of invaded *B. neotomae* (I) or *B. melitensis* (J) by CFU enumeration. (K) *Brucella* invasion assay in iBMDMs treated with CREB and CXCR4 inhibitors. iBMDMs treated with AMD3100 (CXCR4 inhibitor) for 24 hours or CREB inhibitor for 3 hours as indicated, followed by infection with *B. neotomae* and enumeration of CFU for determining the intracellular bacteria. The data shown in A, C, & D panels are representations of three independent experiments. The data shown in B, E, F, G, H, I, J & K panels are representations of two independent experiments. The data are presented as the mean ± SEM (P < 0.05*; P < 0.01**, P <0.001; ***).

Our experimental data suggest that *Brucella* modulates USP8 at the transcriptional level, as mRNA expression levels of USP8 were also affected. Various signalling processes are reported to activate the transcription factor CREB through its phosphorylation that in turn, drives the expression of USP8 (Bland et al., 2019). Therefore, *Brucella* may affect the activation of CREB, which may downregulate the expression of USP8. To examine this, we analysed USP8 levels in macrophages treated with the antagonist of CREB. The iBMDMs were treated with increasing concentrations of CREB antagonist for three hours, followed by qRT-PCR analysis to examine USP8 mRNA level and immunoblotting to assess phospho-CREB and USP8 protein levels. We observed a dose-dependent depletion of the USP8 mRNA level as well as the protein levels of phospho-CREB and USP8 in the cells treated with the CREB antagonist (Figures. 3F & G). To examine whether *Brucell*a infection affects the activation of CREB, the phospho-CREB, and USP8 were evaluated in the infected and uninfected iBMDMs. We observed a significant suppression of CREB phosphorylation and USP8 expression in the *Brucella*-infected macrophages, demonstrating the role of *Brucella* in impacting USP8 transcription by inhibiting the activation of CREB (Figure. 3H).

Next, we analysed whether CREB inhibitor interferes with the invasion of *Brucella* into macrophages. The iBMDMs were treated with the CREB inhibitor for 3 hrs, followed by examining the invasion of *B. neotomae* or *B. melitensis*. We observed a significant enhancement of *B. neotomae* or *B. melitensis* invasion into the cells treated with CREB inhibitor compared to cells treated with the vehicle control, DMSO (Figures. 3I & J). Further, we examined whether the effect of the CXCR4 inhibitor on *Brucella* invasion can be negated by treating cells with the CREB inhibitor. To examine this, iBMDMs were first treated with the CXCR4 inhibitor for 24 hours, followed by treating the cells with the CREB inhibitor for 3 hours and *Brucella* invasion assay. As observed before, CXCR4 treatment resulted in a diminished invasion of *Brucella,* whereas this effect was counteracted by subsequent treatment of CXCR4-treated cells with the CREB inhibitor (Figure. 3K). Collectively, our experimental data suggest that CREB-mediated expression of USP8 affects the levels of CXCR4 that regulate the invasion of *Brucella* into macrophages.

### The *Brucella* effector protein, TcpB, downregulates the expression of USP8

The TLR2/4 adaptor protein, TIRAP, is known to activate the CREB signalling pathway during the inflammatory response in microbial infections (Mellett et al., 2011; Newton and Dixit, 2012). *Brucella* encodes the effector protein, TcpB, which is reported to promote the ubiquitination and degradation of TIRAP to attenuate the TLR2/4 signalling (Alaidarous et al., 2014). Therefore, we sought to examine whether TcpB plays any role in the modulation of USP8 expression. To examine this, we transfected HEK293T cells with increasing concentrations of eukaryotic expression plasmid harbouring TcpB, then analyzed the levels of USP8 by q-RT-PCR and immunoblotting. Interestingly, TcpB suppressed the expression of USP8 in the transfected cells in a dose-dependent manner (Figures. 4A & B). TcpB did not affect the mRNA levels of MYD88 and TIRAP, indicating the specificity of TcpB towards suppressing the USP8 transcription (Figures. S4 A & B). TcpB is a cell-permeable protein, and the recombinant TcpB protein is reported to get translocated into the macrophages (Radhakrishnan and Splitter, 2010). To further confirm the experimental data, iBMDMs were treated with purified MBP or MBP-TcpB protein for 5 hours, then analyzed the level of USP8 expression by immunoblotting. We observed a significant suppression of USP8 in the cells treated with MBP-TcpB compared to MBP alone, confirming the role of TcpB in downregulating the USP8 expression (Figure. 4C). Next, we analysed the levels of phospho-CREB and USP8 in the presence of TcpB. HEK293T cells were transfected with increasing concentrations of HA-TcpB expression plasmid, followed by analysing the endogenous levels of phospho-CREB and USP8 by immunoblotting. The experimental data showed a dose-dependent decrease in phospho-CREB and USP8 levels in the cells transfected with TcpB, confirming the role of TcpB in downregulating the USP8 through suppression of CREB phosphorylation (Figure. 4D).

**Figure 4.**
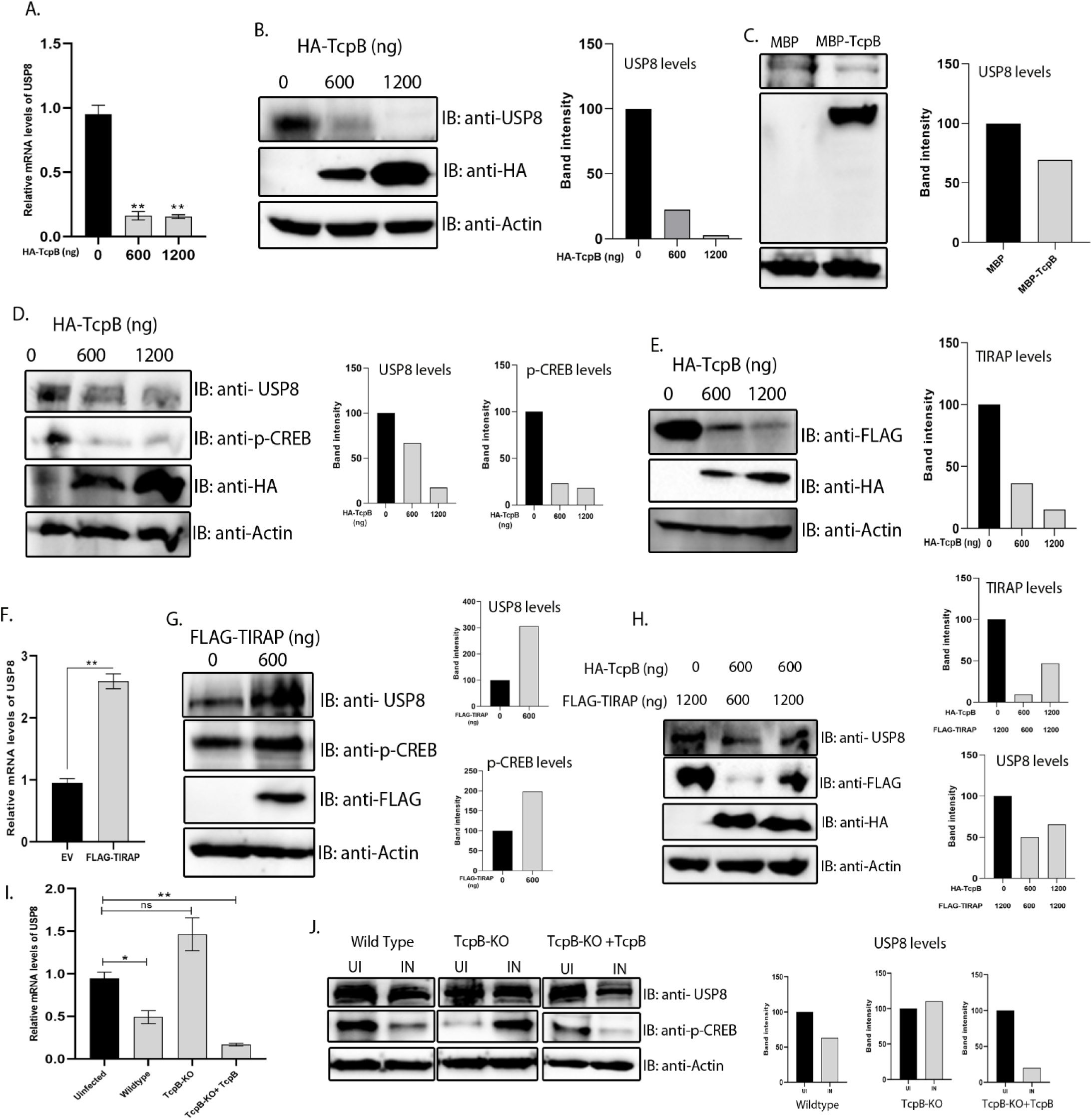
The *Brucella* effector protein, TcpB, suppresses USP8 expression by targeting the TIRAP-CREB signalling pathway. (A) Endogenous levels of USP8 in iBMDMs overexpressing TcpB. The cells were transfected with the indicated concentrations of eukaryotic expression plasmid harbouring HA-tagged TcpB or EV for 24 hours, followed by analysing the mRNA expression level of USP8 by qRT-PCR. The USP8 expression data were normalized with GAPDH, and the relative mRNA expression of USP8 was determined with respect to the cells transfected with EV. (B) Immunoblot showing the levels of USP8 in the HEK293T cells overexpressing TcpB. The cells were transfected with indicated concentrations of eukaryotic expression plasmid harbouring HA-tagged TcpB or EV. Twenty-four hours post-transfection, the cells were lysed and subjected to immunoblotting. The membrane was probed with the anti-USP8 antibody, followed by HRP-conjugated anti-mouse IgG to detect endogenous USP8. HRP-conjugated anti-HA antibody was used to detect HA-TcpB. Actin was used as the loading control. The right panel indicates the densitometry analysis of USP8 bands normalized with actin bands. (C) Immunoblot showing the endogenous USP8 levels in iBMDMs treated with purified recombinant MBP-TcpB or MBP protein. The cells were treated with the purified proteins for 5 hours, followed by lysis and immunoblotting. HRP-conjugated anti-MBP antibody was used for detecting the MBP-TcpB or MBP alone. The right panel indicates the band intensity of USP8, which was normalized with actin bands. (D) Immunoblot showing the endogenous levels of USP8 and phospho-CREB in HEK293T cells transfected with increasing concentrations of the plasmid expressing HA-TcpB. The right panel indicates the densitometry analysis of USP8 and phospho-CREB bands normalized with actin bands. (E) Immunoblot showing the levels of FLAG-TIRAP in HEK293T cells co-transfected with indicated concentrations of eukaryotic expression plasmid harbouring HA-tagged TcpB or EV for 24 hours. The membranes were probed with HRP-conjugated anti-FLAG and anti-HA antibodies to detect FLAG-TIRAP and HA-TcpB, respectively. The right panel indicates the band intensities of FLAG-TIRAP normalized with actin bands. (F) The mRNA expression level of USP8 in the iBMDMs transfected with the expression plasmid harbouring FLAG-TIRAP. Twenty-four hours post-transfection, cells were harvested and subjected to qRT-PCR analysis to quantify the levels of USP8. The USP8 expression data were normalized with GAPDH, and the relative mRNA expression of USP8 was determined with respect to the cells transfected with EV. (G) Immunoblot showing endogenous levels of USP8 and phospho-CREB in HEK293T cells transfected with indicated concentrations of FLAG-TIRAP expression plasmid for 24 hours. The right panel indicates the densitometry analysis of USP8 and p-CREB bands normalized with the actin bands. (H) Immunoblot showing endogenous levels of USP8 in HEK293T cells transfected with indicated combinations of plasmids expressing FLAG-TIRAP and HA-TcpB. The right panel shows the densitometry analysis of USP8 and FLAG-TIRAP bands normalized with actin bands. (I) The mRNA expression levels of USP8 in iBMDMs infected with wild-type or TcpB-KO *B. neotomae* or TcpB-KO *B. neotomae* complemented with the TcpB expression construct. Four hours post-infection, the cells were harvested, followed by quantification of mRNA levels of USP8 by qRT-PCR. The expression of USP8 was normalized with GAPDH, and the relative mRNA expression of USP8 was determined with respect to the uninfected iBMDMs. (J) Immunoblot showing the level of USP8 and p-CREB in iBMDMs infected with wild-type or TcpB-KO *B. neotomae* or TcpB-KO *B. neotomae* complemented with TcpB expression construct. The cells were harvested 4-hours post-infection, followed by immunoblotting to detect endogenous USP8 and phospho-CREB. The right panel indicates the densitometry analysis of USP8 and phospo-CREB bands normalized with actin bands. The data shown in A, B, D, H, I & J panels are the representations of three independent experiments. The data shown in C, E, F & G panels represent two independent experiments. The data are presented as the mean ± SEM (P < 0.05*; P < 0.01**, P <0.001; ***). EV: Empty Vector.

To confirm that TcpB modulates USP8 gene expression through TIRAP, we analysed the levels of TIRAP and phospho-CREB in the TcpB transfected cells. In concordance with the previous reports, we observed a dose-dependent degradation of TIRAP by TcpB in HEK293T cells co-transfected with the eukaryotic expression plasmids harbouring TcpB or TIRAP (Figure. 4E). Further, a dose-dependent degradation of endogenous TIRAP was also observed in HEK293T cells transfected with the increasing concentrations of HA-TcpB expression plasmid (Figure. S4 C). Next; we sought to examine whether TIRAP is required for phosphorylation of CREB which in-turn promotes USP8 transcription. To analyse this, HEK293T cells were transfected with the plasmid harbouring FLAG-TIRAP or empty vector, followed by evaluating the mRNA levels of USP8 by qRT-PCR and endogenous levels of phospho-CREB and USP8 by immunoblotting. As anticipated, we observed an enhanced USP8 transcription and enhanced endogenous phospho-CREB and USP8 levels in the cells that are overexpressing FLAG-TIRAP, indicating the role of TIRAP in activating CREB and its downstream target, USP8 (Figures. 4F & G). To analyse whether overexpression of TIRAP rescues USP8 suppression by TcpB, we performed a titration experiment where HEK293T cells were transfected with 600 ng of TcpB and increasing concentrations of TIRAP (600 and 1200 ng) expression plasmids, followed by evaluating the expression of USP8. We observed a diminished suppression of USP8 by TcpB with the higher concentration of TIRAP, indicating the effect of TcpB on USP8 through TIRAP (Figure. 4H). Collectively, our data show that degradation of TIRAP by TcpB prevents activation of CREB that impacts the transcription of USP8.

To examine the role of TcpB in regulating the USP8 expression further, we generated TcpB knockout (KO) *B. neotomae* and TcpB KO *B. neotomae* complemented with the TcpB expression construct, followed by their confirmation by PCR analysis (Figure. S4 D). Subsequently, the iBMDMs were infected with wild-type *B. neotomae* or TcpB KO *B. neotomae* or TcpB KO *B. neotomae* complemented with TcpB expression plasmid and evaluated the levels of USP8 by qRT-PCR and protein levels of USP8 and phospho-CREB by immunoblotting at 4-hours post-infection. We observed diminished phospho-CREB and USP8 in the cells infected with wild-type, or TcpB KO *B. neotomae* complemented with TcpB expression plasmid. In contrast, phospho-CREB and USP8 levels were unaffected in the iBMDMs infected with TcpB KO *B. neotomae* (Figures. 4I & J). These experimental data indicate that *Brucella* modulates the expression of USP8 through its effector protein, TcpB.

### Inhibition of USP8 enhances the splenic load of *Brucella* in the infected mice

Our *in vitro* studies indicated that inhibition of USP8 resulted in an enhanced invasion of *Brucella* into macrophages through the plasma membrane receptor, CXCR4. Therefore, we wished to examine the effect of USP8 or CXCR4 antagonists *in vivo* using the mice model of brucellosis. Eight-week-old female BALB/c mice were infected with *B. melitensis*, followed by treatment with DUB_IN-2 or BV02 or AMD3100 or vehicle control, 10 days post-infection for 3 days. Subsequently, the mice were sacrificed, and the load of *B. melitensis* in the spleen was estimated by CFU analysis (Figure. 5A). The spleens from the mice treated with DUB_IN-2 showed a significantly higher number of *B. melitensis* than those treated with the vehicle control (Figure. 5B). In contrast, we observed a decreased number of *B. melitensis* in the spleen of mice that were treated with BV02 or AMD3100 as compared to the control mice (Figures. 5C & D). Our *in-vivo* experimental data suggest that the chemical inhibition of USP8 augments the load of *B. melitensis* in mice. In contrast, inhibition of CXCR4 or activation of USP8 leads to a remarkable reduction of *B. melitensis* survival in the infected mice.

**Figure 5.**
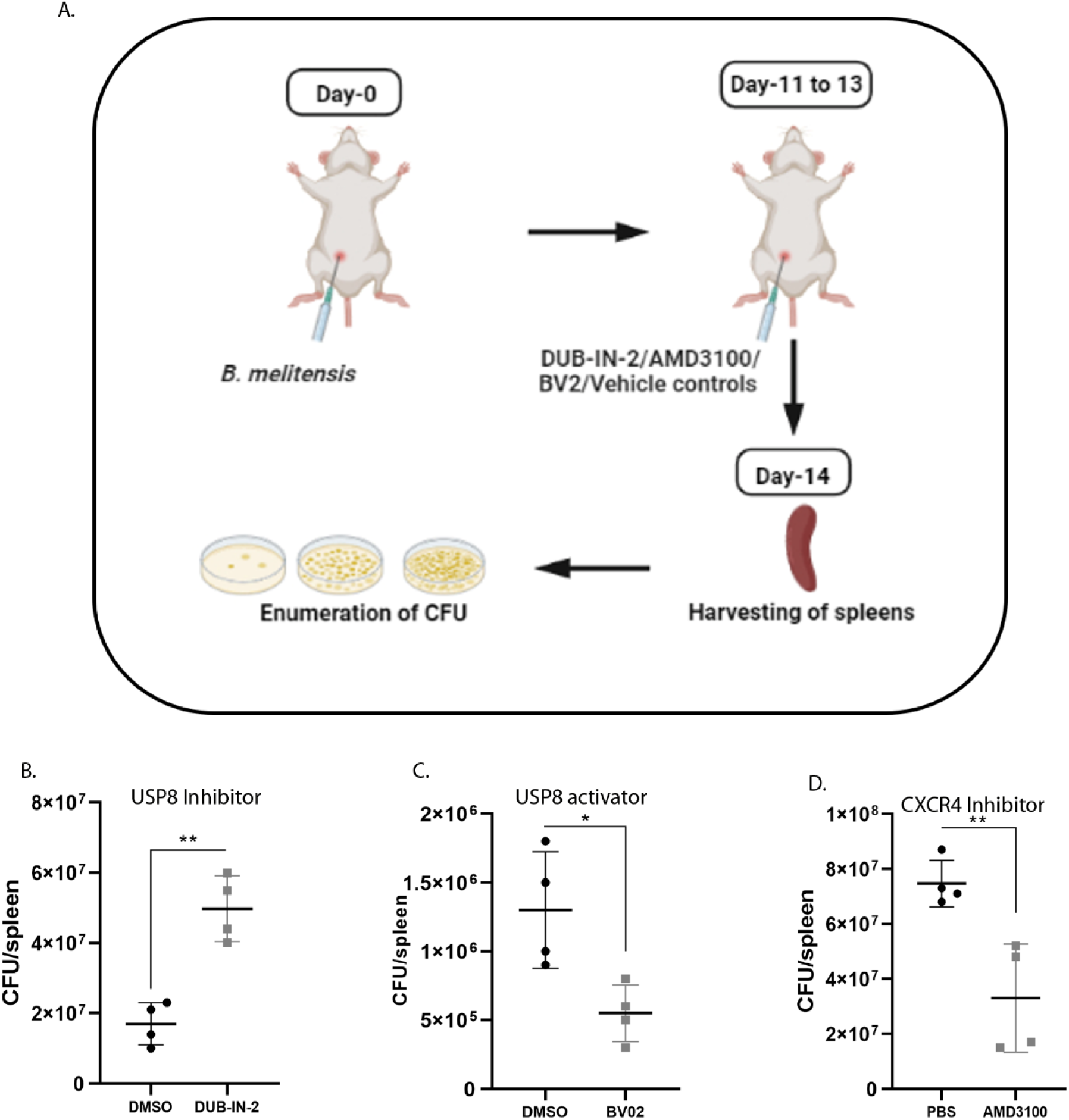
USP8 plays a vital role in determining the load of *Brucella* in the mice infected mice. (A) Schematic diagram showing the methodology used for examining the effect of inhibition of USP8/CXCR4 or activation of USP8 on the load of *B. melitensis* in the infected mice. (B) Scatter plot showing the splenic load of *B. melitensis* in the mice treated with inhibitors of USP8/CXCR4 or activator of USP8. Eight-week-old female BALB/c mice were infected with *B. melitensis*. Ten-day post-infection, infected mice were treated with DUB_IN-2 (B) or BV02 (C) or AMD3100 (D) or vehicle control for 3 days. Subsequently, the mice were sacrificed, and the load of *B. melitensis* in the spleen was estimated by CFU analysis. The data are presented as the mean ± SEM (P < 0.05*; P < 0.01**, P <0.001; ***).

**Figure 6.**
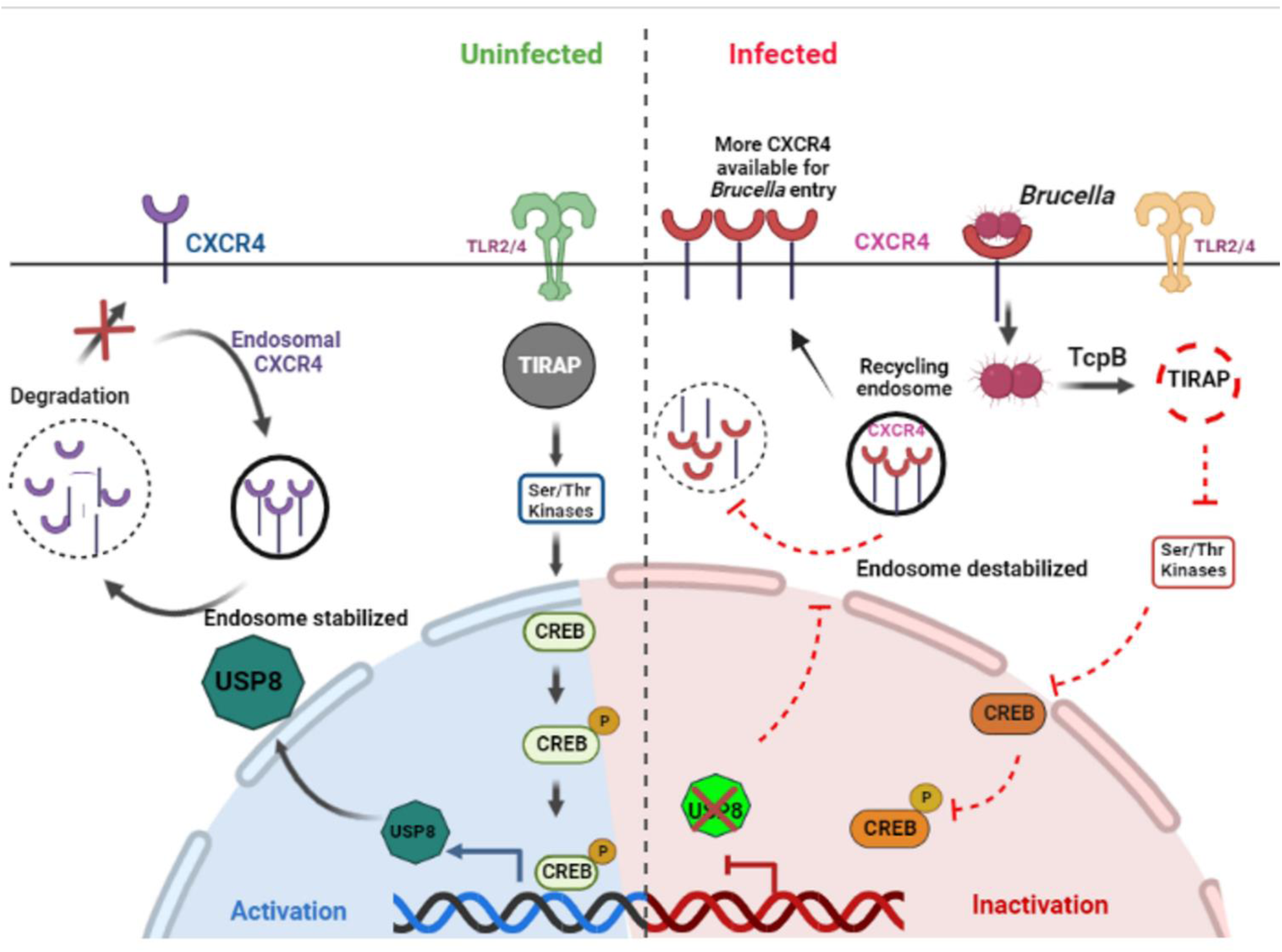
Graphical summary depicting the role of USP8 in regulating the invasion of *Brucella* into the macrophages. Expression of USP8 through the TIRAP-CREB signalling pathway negatively regulates the turnover and plasma membrane localization of the CXCR4 receptor (Left). Upon *Brucella* infection, the effector protein, TcpB, degrades TIRAP, preventing the CREB activation and transcription of USP8 (Right). This promotes the recycling of endosomal CXCR4 and its availability on the plasma membrane, facilitating an enhanced *Brucella* invasion into macrophages.

## Discussion

*Brucella* species successfully invade and multiply in the phagocytic cells by subverting various cellular processes (Głowacka et al., 2018). Their ability to prevent the intracellular killing and to create a safe replication permissive niche in the infected cells contribute to the chronicity of *Brucella* infection. *Brucella* also evades recognition or suppresses the activation of various host immune responses to chronically persist in the host (Jakka et al., 2017; Luo et al., 2017). However, minimal information is available on *Brucella* and host factors involved in host-pathogen interaction. Among many stages of the intracellular cycle of *Brucella*, gaining entry into the host cells is critical for initiating the infection cycle. Here we identified the host protein, USP8, that plays an essential role in the invasion of *Brucella* into macrophages and in determining the bacterial load in the infected host.

USP8 is a cystine protease family of deubiquitinases, which cleaves the conjugated ubiquitin moiety from its protein substrates to regulate various cellular processes (De Ceuninck et al., 2013; MacDonald et al., 2014). It is a multidomain protein containing a microtubule interacting domain, transport domain (MIT), SH3-binding motifs (SH3-BM), 14-3-3-peptide-binding motif, and catalytic domain (DUB). USP8 has pleiotropic functions inside the cells and maintains cellular homeostasis by regulating protein turnover and cargo sorting (Dufner and Knobeloch, 2019). Our studies unravelled a novel function of USP8, where it plays a vital role in the host defence against *Brucella* infection. Downregulation of USP8 expression through siRNA or inhibiting its activity using chemical inhibitors potentiated *Brucella* invasion of macrophages and bacterial load in the infected mice, highlighting its role in the host defence. In contrast, enhancing the USP8 activity compromised the macrophage invasion and bacterial persistence in the infected mice, confirming USP8 as part of the host defence mechanism to avert pathogen entry and survival.

USP8 is reported to regulate the turnover and membrane-localization of the plasma membrane-localized receptor, CXCR4. Though USP8 stabilizes various tyrosine kinase receptors such as EGFR and TβRII through its deubiquitylation property, it has a paradoxical function by enhancing the lysosomal degradation of CXCR4 (Berlin et al., 2010b; Xie et al., 2022). CXCR4 is known to express on various immune cells, osteoclasts, and periodontal tissues. It has been reported to function as a co-receptor for the entry of HIV-1 virion into the host cells (Xiao et al., 2021). The involvement of CXCR4 in the colonization of *Porphyromonas gingivalis* in periodontal tissues through binding to the fimbria has also been reported (McIntosh and Hajishengallis, 2012). The invasion of *Brucella* into macrophages involves the polymerization of actin filaments, receptor-mediated endocytosis, and interaction with various cytoskeletal regulators, such as small GTPases on the plasma membrane (Guzmán-Verri et al., 2001; Watarai et al., 2003). In addition, the CXCR4 receptor is reported to promote the entry of *Brucella* into the macrophages. Our studies revealed that USP8 affects the invasion of *Brucella* into macrophages through modulation of membrane localization and an overall turnover of the CXCR4 receptor. The inhibition of CXCR4 affected the invasion of *Brucella* into macrophages and its survival in the mice, underscoring the essential role of this receptor in invading the host cells. Further, counteracting the effect of USP8 inhibition by blocking CXCR4 confirms the link between USP8 and CXCR4 in the invasion of *Brucella*. The data on mice infection studies also demonstrate the physiological relevance of USP8 suppression upon *Brucella* infection for persistence in the host and the role of membrane receptor, CXCR4 in the entry of *Brucella* into the host cells.

The depletion of CXCR4 through USP8 appears to be an efficient host-defense strategy that operates by negatively regulating the receptors employed by pathogenic microorganisms to gain entry into the cells. However, the pathogens co-evolve with the host, which enables them to successfully overcome the defense mechanism to infect the host. In support of this view, we observed that *Brucella* downregulates the expression of USP8 at its early stages of infection to facilitate an unhindered invasion, which is crucial for its survival in the host. However, the normal expression of USP8 was restored after the invasion part of its life cycle, indicating the precise manipulation of host cellular processes according to the different stages of infection. In agreement with this observation, the upregulation of various scavenger receptors by *Brucella* through the host protein, FBXO22, has also been reported (Mazumdar et al., 2022).

Intriguingly, we observed that only live *Brucella* could impart the suppression of USP8, suggesting the role of a secreted effector protein to hijack the host signalling pathway involved in USP8 transcription. The changes in the expression levels of host genes upon *Brucella* infection are described in various instances previously, signifying the importance of host gene modulations for the *Brucella*-macrophage interaction (von Bargen et al., 2012). Our subsequent experiments identified that the *Brucella* effector protein, TcpB plays an essential role in negatively regulating USP8 expression through the TLR2/4 adaptor protein, TIRAP. Cyclic-AMP Response Element Binding protein (CREB) is a reported USP8 transcription factor that drives the expression of the USP8 gene (Bland et al., 2019). Since CREB acts as a key factor, which regulates downstream signalling events that are triggered during the microbial encounter to activate immune responses, we hypothesized that CREB might be an upstream target of *Brucella* to suppress USP8 gene transcription. Studies have shown that TIRAP-mediated signalling pathways can lead to the activation of CREB through its phosphorylation resulting in the expression of various inflammatory and anti-inflammatory genes during microbial infections (Newton and Dixit, 2012). This indicates that eliminating TIRAP through proteasomal degradation can compromise the CREB activation and expression of CREB-dependent genes. TcpB is known to induce ubiquitination and subsequent degradation of TIRAP to attenuate induction of pro-inflammatory cytokines to subvert the host innate immune responses (Radhakrishnan et al., 2009). Our subsequent studies revealed that TcpB-induced TIRAP degradation prevents CREB activation and transcription of USP8. Infection studies with TcpB KO *Brucella* further confirmed the role of TcpB in *Brucella*-mediated suppression of USP8 through the CREB signalling pathway. Studies have shown that *Brucella* employs its effector proteins to manipulate the host cellular pathways to create a replication permissive niche in the infected cells. The functional interaction of *Brucella* effectors, BspB and RicA is reported to mediate Rab2-dependent transport in the intracellular life cycle of *Brucella* (Smith et al., 2020). Two recently identified *Brucell*a effector proteins, NyxA and NyxB, are shown to modulate SENP3 affecting the subcellular localization of nucleolar proteins (Louche et al., 2023).

In conclusion, our experimental data unravel a previously unknown role of USP8 as an essential player in the host defence against microbial infections. USP8 effectively prevents the invasion of *Brucella* into macrophages by depleting the membrane receptor CXCR4. Since *Brucella* species live in close association with the mammalian hosts, they have evolved efficient ways to invade the host cells. In support of this view, we found that *Brucella* negatively regulates the expression of USP8 at the early stages of the infection process using its effector protein, TcpB. TcpB exerts its effects through targeted degradation of TIRAP, thereby preventing the activation of CREB and attenuating the USP8 expression. Our findings delineate the mechanisms the host employs to defend against microbial infection and the subversion of these innate defence strategies by the pathogen to establish the infection. Further, the enhancement of the splenic load of *Brucella* upon USP8 inhibition and diminished *Brucella* survival upon activation of USP8 in mice underscores the role of USP8 in determining the host responses to the infection. This information can be extrapolated to other invasive infectious pathogens to understand their mode of infection and persistence in the host. In addition, the various proteins and cellular pathways addressed in this study may serve as potential targets for developing novel therapeutics for brucellosis.

## Author Contribution

Conceptualization, resources, supervision and finding acquisition: GKR; Methodology: GKR and KJ; Investigation: KJ and VBM; Writing–original draft: GKR and KJ; Writing–Review & Editing: GKR and KJ.

